# 3D Convolutional Neural Networks for Classification of Alzheimer’s and Parkinson’s Disease with T1-Weighted Brain MRI

**DOI:** 10.1101/2021.07.26.453903

**Authors:** Nikhil J. Dhinagar, Sophia I. Thomopoulos, Conor Owens-Walton, Dimitris Stripelis, Jose Luis Ambite, Greg Ver Steeg, Daniel Weintraub, Philip Cook, Corey McMillan, Paul M. Thompson

## Abstract

Parkinson’s disease (PD) and Alzheimer’s disease (AD) are progressive neurodegenerative disorders that affect millions of people worldwide. In this work, we propose a deep learning approach to classify these diseases based on 3D T1-weighted brain MRI. We analyzed several datasets including the Parkinson’s Progression Markers Initiative (PPMI), an independent dataset from the University of Pennsylvania School of Medicine (UPenn), the Alzheimer’s Disease Neuroimaging Initiative (ADNI), and the Open Access Series of Imaging Studies (OASIS) dataset. The UPenn and OASIS datasets were used as independent test sets to evaluate the model performance during inference. We also implemented a random forest classifier as a baseline model by extracting key radiomics features from the same T1-weighted MRI scans. The proposed 3D convolutional neural network (CNN) model was trained from scratch for the classification tasks. For AD classification, the 3D CNN model achieved an ROC-AUC of 0.878 on the ADNI test set and an average ROC-AUC of 0.789 on the OASIS dataset. For PD classification, the proposed 3D CNN model achieved an ROC-AUC of 0.667 on the PPMI test set and an average ROC-AUC of 0.743 on the UPenn dataset. Model performance was largely maintained when using only 25% of the training dataset. The 3D CNN outperformed the random forest classifier for both the PD and AD tasks. The 3D CNN also generalized better on unseen MRI data from different imaging centers. These approaches show promise for screening of PD and AD patients using only T1-weighted brain MRI, which is relatively widely available. This model with additional validation could also be used to help differentiate between challenging cases of AD and PD when they present with similarly subtle motor and non-motor symptoms.

## 1. INTRODUCTION

Alzheimer’s disease (AD) and Parkinson’s disease (PD) are the two most common neurodegenerative diseases in the world. According to the World Health Organization (WHO), around 50 million people have dementia worldwide, and AD likely contributes to 60-70% of these cases.^1^ A further study reported that 6.1 million people were affected by PD globally in 2016.^2^ Structural magnetic resonance imaging (MRI) offers the potential to screen individuals for diagnostic biomarkers of PD and AD. Standard volumetric T1-weighted brain MRI is widely used and accessible world-wide, and has the advantage of not subjecting the patient to ionizing radiation^3^ compared to other modalities, such as PET.

Recently, a multi-center study was conducted by the PD working group of the Enhancing NeuroImaging Genetics through Meta-Analysis (ENIGMA) consortium on a cohort of 2,357 PD patients and 1,182 healthy controls using T1-weighted brain MRI scans from 19 different international centers.^4^ The authors analyzed cortical thickness, regional surface area, and volumes for subcortical regions of the brain revealing a distributed pattern of atrophy that advances as the disease progresses. Similar work in AD has stemmed from large cohort studies such as the Alzheimer’s Disease Neuroimaging Initiative (ADNI), which has published multiple works demonstrating significant changes in brain metrics in the disorder.^5^ While important, such case-control comparisons only identify distinctive features at the group level, and do not offer a means to classify disease status in new individuals. Over 80% of PD patients develop cognitive deficits; this is relatively common in prodromal and *de novo* PD. Therefore, differentiating early AD from early PD is a key clinical issue.

Machine learning and deep learning models have been developed for various neuroimaging applications, including image reconstruction and enhancement, segmentation and labeling of regions of interest, and disease classification and subtyping. When designing models for these tasks, several factors must be considered, including data availability, the size and efficiency of the neural network architecture, and its performance on diverse datasets other than those used to train the algorithm. Many studies have shown promise in applying convolutional neural networks (CNNs) to brain MRI scans for regression tasks such as brain age prediction, a commonly used benchmarking task that attempts to predict a person’s age from their brain scan.^6,7^ Other works have proposed methods for classification of neurodegenerative diseases. Islam and Zhang proposed a 2D model based on the Inception-V4 architecture for classification of AD.^8^ Sivaranjini and Sujatha^9^ proposed a CNN based on the AlexNet architecture for PD classification. Many of the methods proposed previously, however, make predictions in a slice-wise manner. Yagis et al. address potential data leakage issues in some of the earlier neuroimaging works, several of which performed slice-based predictions.^10^ Also, some of the prior 2D methods may not account for the spatial relationships of all the slices in the 3D MRI volume. West et al. show that 3D CNNs can outperform their 2D counterparts for some neuro-imaging based classification tasks.^3^

In a recent innovation, Lu et al.^11^ pooled over 85,000 brain MRI scans to build a CNN that classified a person’s sex from their brain scan, and they fine-tuned the network for AD classification using transfer learning. In our own prior work,^12,13^ we used a recurrent neural network (RNN) to aggregate information from an ordered series of 2D slice-based CNNs to predict a person’s age from their 3D MRI scan (see also Peng et al.^14^). We later extended the same method using attention modules to perform AD classification.^6^ Despite promising performance, we did not test the method on additional cohorts beyond the one used to train the algorithm, nor did we test the method on multiple diseases, which perhaps represents a more clinically relevant application.

### Contributions

In this paper, we propose a 3D CNN that extracts features from 3D T1-weighted brain MRI scans and classifies subject-level PD and AD. The key contribution of this work includes:

- A 3D CNN architecture for classification of PD and AD versus healthy control subjects using T1-weighted volumetric brain MRI.
- We test a random forest-based classifier with well-known radiomics features as a baseline model. The 3D CNN outperforms the baseline and generalizes better.
- We evaluate our models on multiple datasets, including external datasets from different centers to independently test our proposed model.
- We show that model performs the classification task reasonably well when trained on a subset of the training data.

## 2. DATA

### 2.1 Datasets

The datasets used in this work include 3D volumetric brain MRI data from the Parkinson’s Progression Markers Initiative (PPMI), University of Pennsylvania (UPenn) as the out-of-distribution test set. PPMI collects baseline scans for untreated patients, and the UPenn is a convenience cohort of treated PD patients. For classification of AD, we analyzed non-accelerated 3D T1-weighted MRI scans from the ADNI dataset and the Open Access Series of Imaging Studies (OASIS) dataset based on OASIS1 (here after referred to as OASIS). Demographics of each dataset are summarized in **Table 1**.

**Table 1.** Summary of the datasets we analyzed in this paper (Parkinson’s Progression Markers Initiative (PPMI), University of Pennsylvania (UPenn), Alzheimer’s Disease Neuroimaging Initiative (ADNI), Open Access Series of Imaging Studies (OASIS))

| Dataset | Total Participants (n) | Controls | Cases (PD or AD) | Sex (Male/Female) | Age Range (years) | Average Age ± Std (years) |
| --- | --- | --- | --- | --- | --- | --- |
| PPMI | 471 | 130 | 341 (PD) | 308M/163F | 30.6-84.9 | 61.10±10.04 |
| UPenn | 117 | 13 | 104 (PD) | 75M/42F | 50.0-84.0 | 66.65±8.12 |
| ADNI | 531 | 243 | 288 (AD) | 268M/263F | 55.7–91.4 | 73.88±7.10 |
| OASIS | 234 | 188 | 46 (AD) | 78M/156F | 33.0-96.0 | 72.38±12.08 |

The H&Y staging characteristics of subjects in the UPenn dataset were as follows: Stage 1 (n=8; 7.1%), Stage 2 (n=36; 32%), Stage 3 (n=64; 56.6%), Stage 4 (n=3; 2.6%), and Stage 5 (n=2; 1.7%). The H&Y staging characteristics of subjects in the PPMI dataset were as follows: Stage 1 (n=151; 44%), Stage 2 (n=191; 55.7%), Stage 3 (n=1; 0.3%), Stage 4 (n=0; 0%), and Stage 5 (n=0; 0%). This H & Y staging information is visualized in **Figure 1** below.

**Figure 1.**
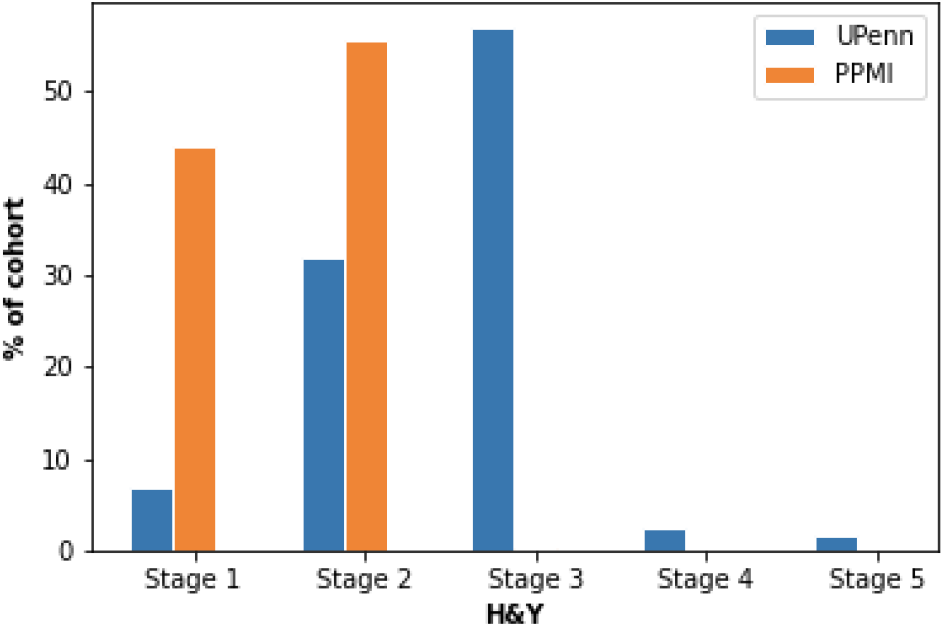
Visualization of H & Y staging information for PPMI and UPenn datasets.

### 2.2 Image Pre-processing

The T1-weighted brain MRI volumes were pre-processed^15^ using a sequence of steps, First we used the fslreorient2std (FSL v6.0.1) to reorient images to match the orientation of the standard template images. We then performed a ‘skull-stripping’ brain extraction using the HD-BET CPU implementation.^16^ Next we extracted grey- and white-matter masks using FSL-FAST (FSL v6.0.1 Automated Segmentation Tool). We then performed a nonparametric intensity normalization (N4 bias field correction)^17^ using ANTs (v2.2). Next we performed linear registration to a UK Biobank minimum deformation template using FSL-FLIRT (FSL v6.0.1 Linear Image Registration Tool) with 6 degrees of freedom. We then performed isometric voxel resampling to 2 mm using the ANTs *ResampleImage* tool (v2.2)^18^, leaving us with pre-processed images of size 91 x 109 x 91. The T1-weighted MR images were scaled to take values between 0 and 1 using a min-max scaling approach for the 3D CNN model.

### 2.2 Data Preparation

The sequence of steps in our preprocessing pipeline ensures some level of uniformity in our datasets, including: nonparametric intensity normalization (N4 bias field correction), and isometric voxel resampling to 2 mm. We created non-overlapping data partitions to train, validate and test the proposed model. 10% of PPMI (PD) and 10% of ADNI (AD) were used for testing for the classification tasks. As there are more PD cases in the UPenn dataset than healthy controls and more controls in OASIS than AD cases, we created balanced subsets of the UPenn (PD) and OASIS (AD) datasets for independent testing. Balanced test data sets for these two cohorts were created by combining the class with fewer subjects with random unique equal sized subsets from the larger classes. Any variance in the performance over the subsets is presented with a corresponding standard deviation in the Results section. Creating these balance subsets allows us to make complete use of all the subjects we have available for classifying PD and AD. We used the rest of the ADNI and PPMI dataset for training (including 20% of these datasets as a validation set for the 3D CNN).

## 3. METHODS

### 3.1 Random Forest Classifier

We implemented a machine learning pipeline as a baseline for AD and PD classification, as shown in **Figure 2**. The model uses the T1-weighted MRI volume and a whole-brain mask as the input. The whole brain mask is created to focus feature extraction on the region within the brain volume. 93 radiomics features - including textural features based on first-order statistical co-occurrence matrices - were extracted using PyRadiomics.^19^ The features were selected based on recursive feature elimination, and were min-max scaled, and used to train a random forest classifier. The random forest was initialized and trained with the default hyperparameters as defined in the *scikit-learn* package. The final model evaluated on the test set is model trained on the complete training dataset.

**Figure 2.**
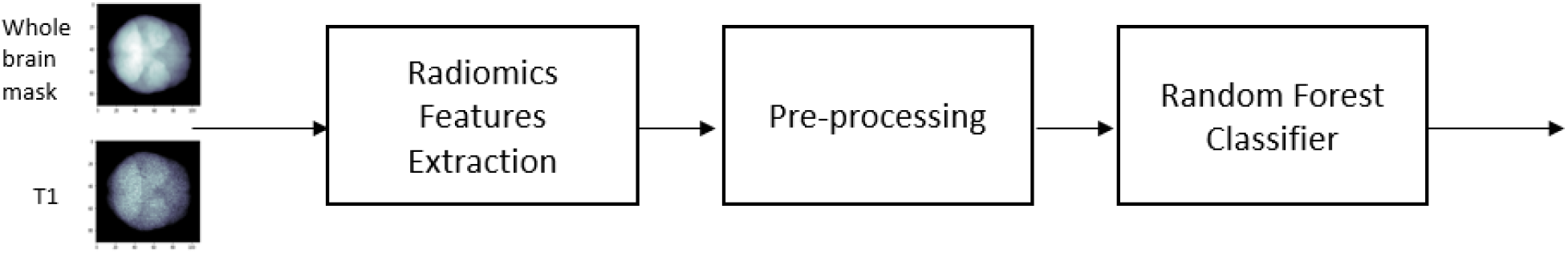
Machine Learning Pipeline for Neurodegenerative Disease Prediction

### 3.2 3D CNN Model Architecture

The architecture of the 3D CNN model^20^ is illustrated in **Figure 3**. The input to this model is 91×109×91. The CNN consists of 4 modules consisting of 3 layers: convolutional (*conv*), max-pooling (*maxpool*) and batch normalization (*BN*) for feature extraction. The convolutional layers have 64, 64, 128, and 256 filters respectively. The feature extractor block is connected by a global average pooling layer to a fully connected (*FC*) layer with 512 neurons, a dropout layer and a fully connected layer with a sigmoid activation function for the binary classification problem. Binary cross-entropy loss is used as the loss function for the CNN.

**Figure 3.**
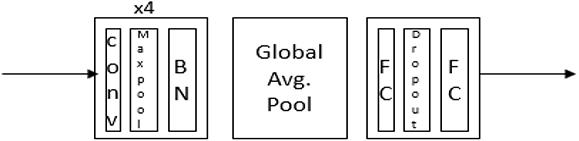
Architecture of the 3D Convolutional Neural Network (CNN).

### 3.3 3D CNN Model Training and Hyperparameter Optimization

All layers of the CNN were trained on the PPMI and ADNI training data for PD and AD classification, respectively. The ROC-AUC and the loss were used to ensure convergence during each of the training epochs. The hyperparameters of the CNN were optimized based on a random search approach. The key hyperparameters were tuned included the optimizer (Adam,^21^ Adam with weight decay, stochastic gradient descent), learning rate (2e-3 to 1e-5), learning rate scheduling, dropout (0 to 0.5), use of bias initializer for class imbalance, early stopping, training epochs, and batch size.

The hyperparameters used for classification of AD included training for 50 epochs, dropout of 0.2, batch size of 4, an Adam optimizer with learning rate of 2e-4, and a weight decay of 1e-4. The hyperparameters used for classification of PD included training for 50 epochs with early stopping if there was no improvement in validation loss for 30 epochs, dropout of 0.2, batch size of 4, an Adam optimizer with learning rate of 2e-4, and weight decay of 1e-4. We used small random image rotations with angles ε {−20, −10, −5, 5, 10, 20} degrees to augment the existing data. We initialized a bias in the final output layer equal to the logarithm of the number of samples in class 0 divided by the number of samples in class 1.^22, 23^ This was performed to alleviate the imbalance in the PPMI dataset. The model with best validation performance was used for testing on the test sets for the two classification tasks.

### 3.4 Model Testing

To evaluate the models, we used standard performance metrics including the receiver-operator characteristic curve-area under the curve (ROC-AUC), the precision-recall area under the curve (PR-AUC), accuracy, precision, recall and F1-score. The Youden index^24^ was used to optimize the threshold from the ROC curve. The performance metrics were calculated as an average on all the balanced subsets of the UPenn and OASIS independent test sets.

## 4. RESULTS

Below we show results of the proposed 3D CNN for classification of PD and AD with T1-weighted MRI data.

### 4.1 Alzheimer’s Disease Classification

The training, validation ROC-AUC, and classification loss curves are shown in **Figure 4** for the 3D CNN. The validation curves closely follow the training performance as seen in the figure. The ROC curves for the 3D CNN, calculated from these test sets, are shown in **Figure 5**. The curves in different colors in the plot on the right in **Figure 5** represent the ROC curves for the balanced subsets from the corresponding test set.

**Figure 4.**
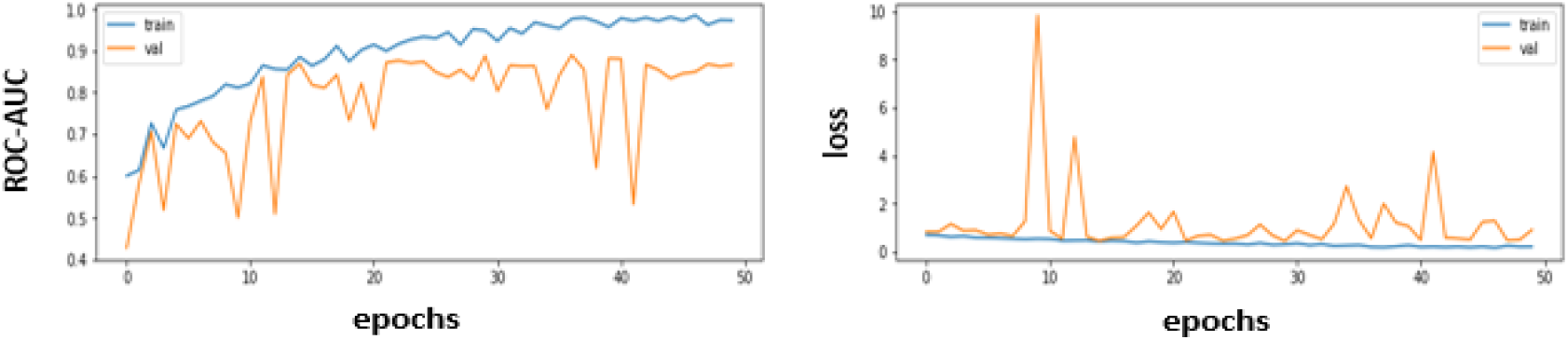
Training and validation curves for ROC-AUC (*left*) and loss (*right*) for AD classification using the 3D CNN.

**Figure 5.**
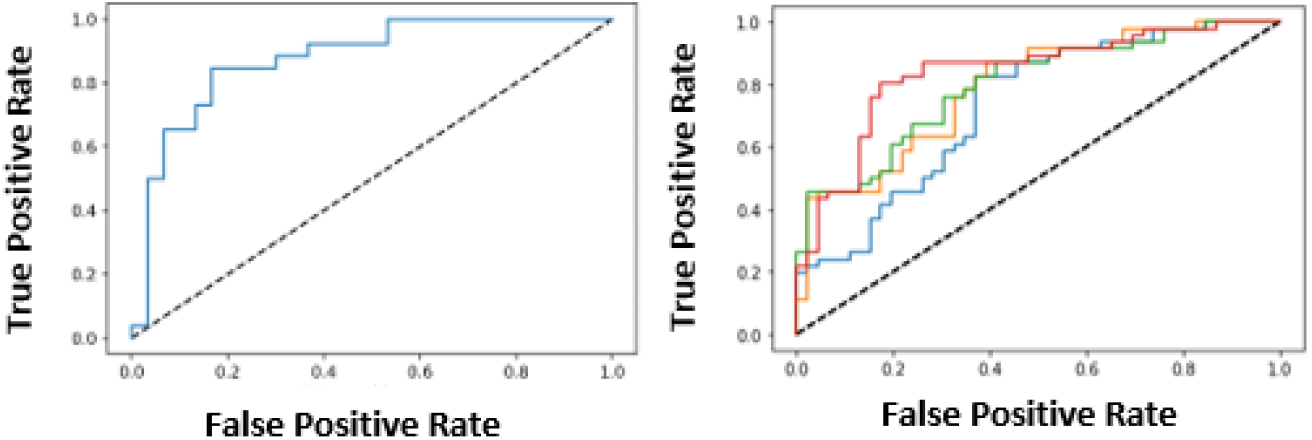
ROC curve for the 3D CNN on the ADNI test set (*left*) and ROC curves for each of the balanced subsets from the OASIS dataset (*right*). The ROC curves for these subsets are shown in different colors on the right

The random forest classifier achieved an ROC-AUC of 0.730 on the ADNI test set and 0.558 on the OASIS independent test set. The 3D CNN achieved an ROC-AUC of 0.878 on the ADNI test set and 0.789 on the OASIS independent test set. The performance of the 3D CNN and random forest classifier on the ADNI test set and the OASIS dataset are summarized in terms of the different metrics in **Table 2**. Results show that our 3D CNN outperforms the baseline random forest for AD classification and tends to generalize better on an independent dataset.

**Table 2.** Performance of Random Forest and 3D CNN for AD classification.

| Metric | ADNI |  | OASIS<br>(Average of balanced subsets with the standard deviation) |  |
| --- | --- | --- | --- | --- |
|  | Random Forest | 3D CNN | Random Forest | 3D CNN |
| ROC- AUC | 0.730 | **0.878** | 0.558 (0.038) | **0.789 (0.038)** |
| PR AUC | 0.646 | **0.816** | 0.556 (0.048) | **0.795 (0.042)** |
| Accuracy | 0.741 | **0.821** | 0.571 (0.018) | **0.742 (0.036)** |
| Precision | 0.722 | **0.808** | 0.575 (0.041) | **0.724 (0.055)** |
| Recall | 0.591 | **0.808** | 0.603 (0.105) | **0.793 (0.039)** |
| F1-score | 0.650 | **0.808** | 0.580 (0.037) | **0.755 (0.029)** |

We evaluated the effect of training data size on the test time performance of the model using ADNI and OASIS. To this end, we used different fractions of the ADNI dataset for training, ranging from just 10% to 100%. We also evaluated the effect of using data augmentation during training. **Figure 6** shows the model performance trend for ADNI and OASIS with and without data augmentation.

**Figure 6.**
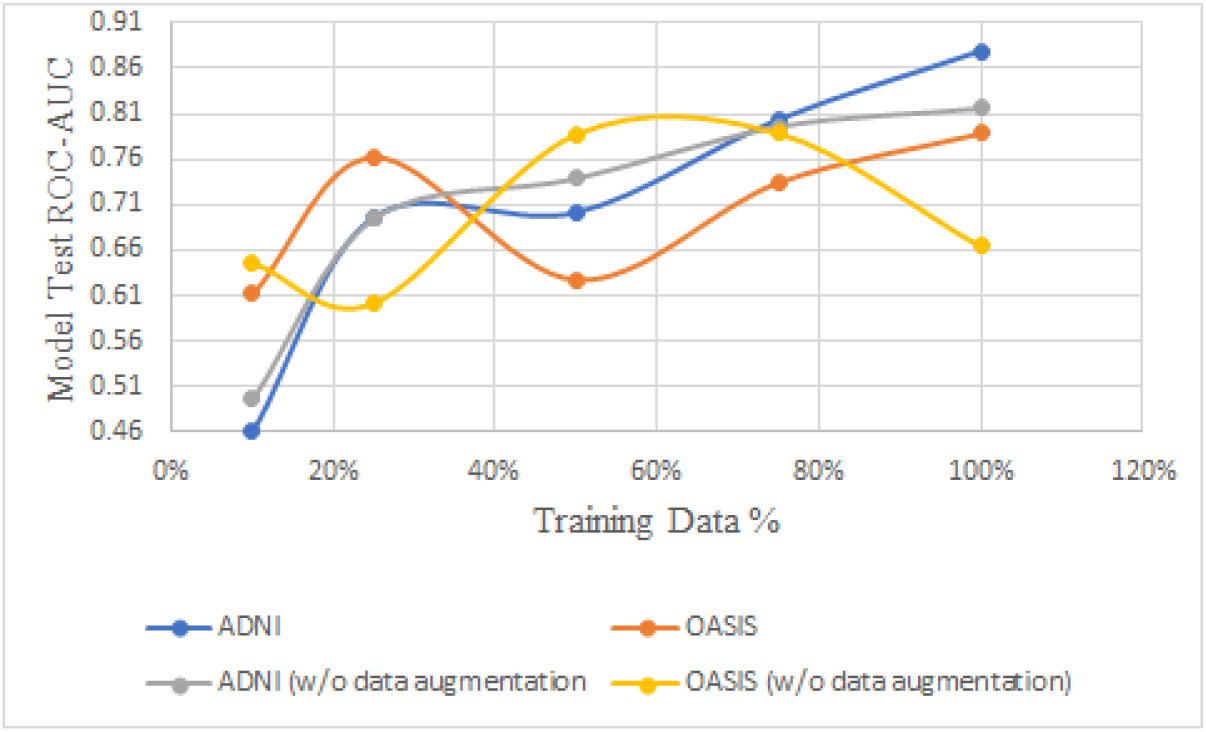
Model test ROC-AUC (for the ADNI test data and for OASIS) is plotted versus the Training Dataset size (from the ADNI dataset), On the y-axis we show performance for models trained on 10% (n=37), 25% (n=92), 50% (n=185), 75% (n=278), and 100% (n=371) of the available training data.

### 4.2 Parkinson’s Disease Classification

The training, validation ROC-AUC and classification loss curves are shown in **Figure 7** for the 3D CNN for PD classification. **Figure 8** shows the ROC curves for the 3D CNN on the PPMI test set, and over each balanced subset of the UPenn dataset.

**Figure 7.**
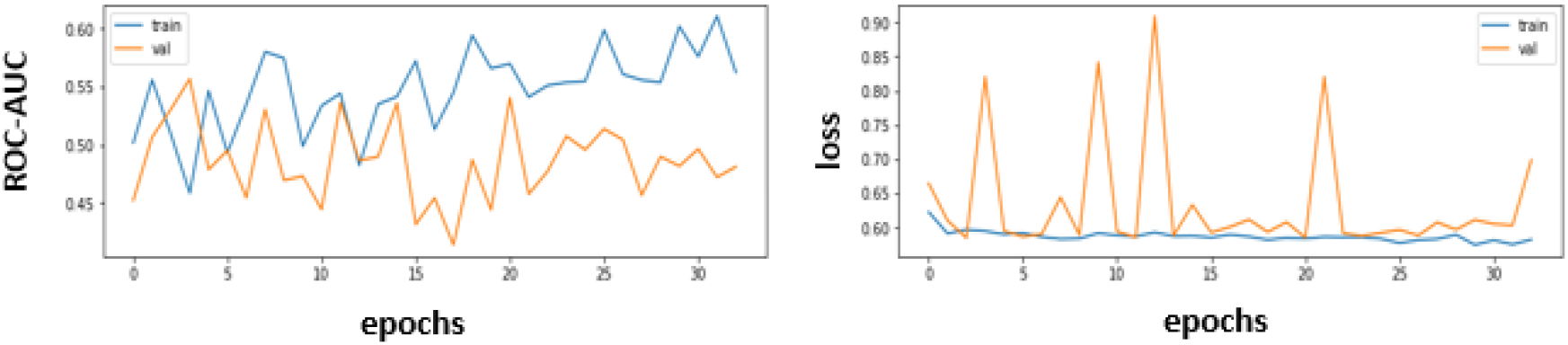
Training and validation curves for ROC-AUC (*left*) and loss (*right*) for PD classification using the 3D CNN.

**Figure 8.**
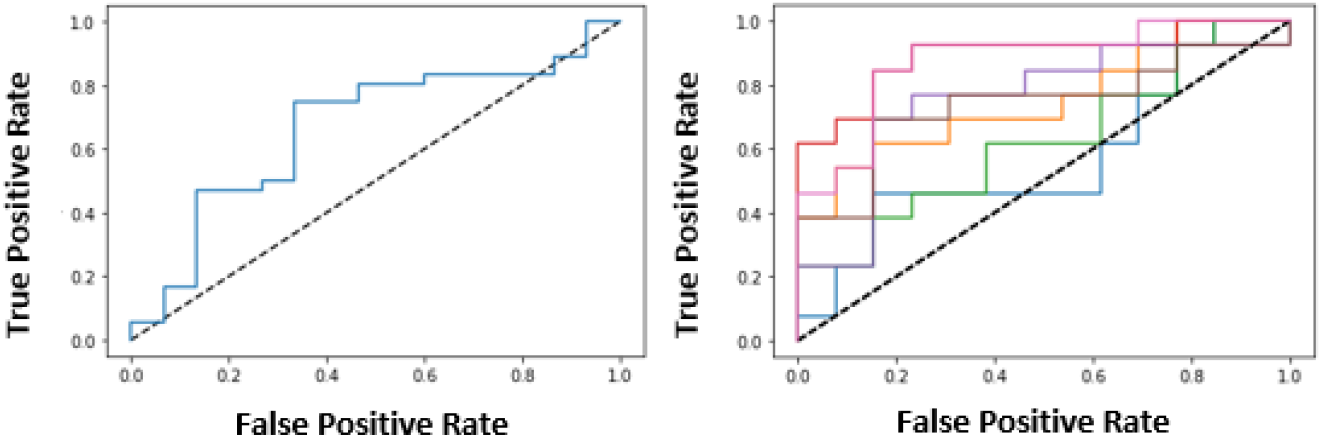
ROC curve of the 3D CNN on the PPMI test set (*left*) and ROC curves for each of the balanced subsets from the UPenn dataset (*right*). The ROC curves for these subsets are shown in different colors on the right.

The random forest classifier achieved an ROC-AUC of 0.524 on the PPMI test set and 0.534 on the UPenn test. The 3D CNN achieved an ROC-AUC of 0.667 on the PPMI test set and 0.743 on the UPenn test. The performance of the 3D CNN and random forest classifier during inference on the PPMI test set and the UPenn independent test set are summarized using the different metrics in **Table 3**. Results show that our 3D CNN outperforms the baseline random forest for PD classification and generalizes better when tested on an independent dataset.

**Table 3.** Performance of Random Forest and 3D CNN for PD classification

| Metric | PPMI |  | UPenn<br>(Average of balanced subsets with the standard deviation) |  |
| --- | --- | --- | --- | --- |
|  | Random Forest | 3D CNN | Random Forest | 3D CNN |
| ROC AUC | 0.524 | **0.667** | 0.534 (0.066) | **0.743 (0.111)** |
| PR AUC | 0.815 | **0.812** | 0.520 (0.067) | **0.777 (0.102)** |
| Accuracy | 0.521 | **0.706** | 0.567 (0.066) | **0.709 (0.082)** |
| Precision | 0.864 | **0.839** | 0.563 (0.071) | **0.816 (0.085)** |
| Recall | 0.487 | **0.722** | 0.673 (0.248) | **0.560 (0.208)** |
| F1-score | 0.623 | **0.776** | 0.581 (0.148) | **0.631 (0.177)** |

## 5. DISCUSSION AND FUTURE WORK

In this paper, we present a 3D CNN that outperforms a baseline random forest method for the classification of AD and PD from T1-weighted volumetric brain MRI. We presented results comparing the 3D CNN’s performance with the baseline method, for both tasks, in section 4. The 3D CNN achieved an ROC-AUC of 0.878 and 0.667 for the AD (10% of ADNI) and PD (10% of PPMI) classification. The 3D CNN yielded an average ROC-AUC of 0.789 and 0.784 for the AD (OASIS) and PD (UPenn) classification. The standard deviation on the UPenn test (0.11) for the 3D CNN versus on the OASIS test set (0.03) suggests that the model tuned for the AD task is relatively more stable than on the PD task. This can also be observed via the ROC curves on the right of **Figures 5** and **8**. The random forest classifier achieved an ROC-AUC of 0.730 and 0.524 for the AD (10% of ADNI) and PD (10% of PPMI) classification. The random forest yielded an average ROC-AUC of 0.558 and 0.534 for the AD (OASIS) and PD (UPenn) classifications. The 3D CNN also generalized better than the random forest classifier on the independent test sets.

We show how the size of the training dataset affects the model’s performance during inference. As expected, the model performance degrades with a smaller training dataset. Even so, the model performed reasonably well with only 25% of the ADNI training data, i.e., 92 subjects. **Figure 6** indicates that data augmentation could be beneficial to the model’s performance by reducing overfitting during training. The performance trend shows that the model benefits from using additional training data and may continue to improve as more training datasets become available.

Empirically, the 3D CNN was more prone to overfitting on the PPMI data for PD classification than on ADNI for AD classification. PD classification might be more challenging than AD classification, as definitive diagnosis of PD might require other imaging modalities such as DAT-PET to augment structural features from T1-weighted brain MRI. Future work will also assess the added value of additional imaging protocols, such as quantitative MRI,^25^ which may boost classification performance. Our PPMI dataset was imbalanced for the two classes, i.e., the class ratio was 0.72(PD):0.28(CN), whereas ADNI was relatively well balanced, i.e., 0.46(AD):0.54(CN). A larger dataset with more balance between the classes might improve performance and reduce overfitting during training for PD classification. The 3D CNN proposed here is a relatively efficient architecture (~1.3 million parameters) compared to other typical 3D deep neural networks. We plan to perform additional experiments with 2D variants as this might also help to reduce overfitting issues for low data problems such as the tasks defined here. We are also working on creating activation maps as a way to improve interpretability of the model’s prediction, and plan to test the added value of MRI data harmonization approaches based on generative adversarial networks, such as CycleGANs^26^ or CALAMITI,^27^ which can make MRI data from different scanners or protocols more comparable.

## 6. CONCLUSIONS

The experiments and results presented here highlight the potential of the 3D CNN model to classify neurodegenerative diseases based on T1-weighted brain MRI. The 3D CNN’s performance was robust when trained only on a subset of the complete dataset, and outperformed the baseline random forest classifier. We showed some differences in training and test performance for the proposed model in PD and AD classification tasks. When trained on more data and with additional validation the proposed pipeline could help in non-invasively screening individuals for PD and AD from structural MRI data. Even classifiers with moderate performance may be useful screening tools, prior to more invasive procedures, such as PET scans and CSF assays, which have been used to define biological AD and to help diagnose PD.

## Acknowledgements

This work was supported by in part by the U.S. National Institutes of Health, under NIH grants U19 AG062418, R01 NS107513, P01 AG066597 (Imaging Core), U01 AG068057, by DARPA under Agreement HR00112090104, and by a Zenith Award from the U.S. Alzheimer’s Association.

